# Computational Modeling of Hyperpolarizing Astrocytic Influence on Cortical Up-Down State Transitions

**DOI:** 10.1101/2023.10.16.562461

**Authors:** Jay Verma, Pranjal Garg

## Abstract

The Up-Down dynamics seen in cortical structures during non-rapid-eye-movement (NREM) sleep, anesthetized states, and quiet wakefulness is the spontaneous alternation between phases of heightened firing activity (referred to as the Up state) and periods of neuronal inactivity (termed the Down state) within neural networks. By leveraging bistable dynamics imposed by a depolarising astrocyte population, in the current paper, we introduced a hyperpolarising astrocyte population to an existing model of Up-Down dynamics to account for biological relevance. We created a computational rate model that includes populations of depolarizing and hyperpolarizing astrocytes and neurons. To optimize model parameters, we used the Elementary Effects (EE) test. It was followed by linear stability analysis to locate bistable regimes in the parameter hyperspace. The addition of hyperpolarizing gliotransmission perturbed model dynamics, indicating its sensitivity to qualitatively differing architectures. We then identified a bistable regime within the dynamics spectrum. According to the EE test, the strength of cell population coupling is a low-sensitivity parameter, possibly due to neuroplastic changes. We also found that the threshold of excitatory cell populations and the strength of adaptation are high-sensitivity parameters, whereas the threshold of inhibitory cell populations is low-sensitivity. Our model enables the possibility of testing biologically relevant theories of hyperpolarizing gliotransmission, where data remains scant.

## 1 Introduction

Emergent behaviors arise from the network properties of neural systems. Understanding the mechanisms and properties of such behaviors frequently requires a combination of experimental and theoretical approaches. One of these behaviors that are actively being studied is the cortical Up-Down dynamics, characterized by periods of neuronal firing alternating with periods of silence, even in the absence of external inputs, and spontaneous transitions between these two states. This pattern of cortical activity is usually observed during non-rapid-eye-movement sleep (NREM), anesthetized states, and quiet wakefulness (1-4).

The cellular mechanisms responsible for generating Up-Down dynamics are an area of active research. Several theoretical studies have explored models of Up-Down dynamics, incorporating varying degrees of biological complexity and plausibility (5-8). These models generally comprise a neuronal network with an embedded negative feedback mechanism that allows the system to alternate between the two states (7-11). A key experimental observation is that the Up-Down pattern appears to be intrinsic to the cerebral cortex, not to imply that external inputs, such as those from the thalamus and the limbic system, can’t influence it (3, 12-14). However, this finding makes the task of constructing an intuitive yet plausible model much more conceivable.

An ever-increasing body of evidence is being generated on the diverse roles of astrocytes in the nervous system. Classically, they were considered connective tissue cells that are a component of the glia. In contrast, recent research shows that astrocytes can also act as integrators of neuronal electrical activity and form a tripartite synapse that exhibits multidirectional information exchange among the presynaptic and postsynaptic neurons, and the astrocyte (15-18). This has been accompanied by concurrent advances in theoretical literature that acknowledge these complex roles of astrocytes, as exemplified by Moyse and Berry, 2022 (19).

Building upon the work of Jercog, Roxin, et al., 2017 (20), Moyse and Berry, 2022 described a relatively simple model that exhibited bistability and spontaneous transitions between the Up and Down states. One of the key conclusions from their work is that astrocytes can indeed be hypothesized to participate in synaptic information exchange via gliotransmitters. Therefore, it is an argument in favor of the importance of gliotransmission, which is a subject of intense debate among neuroscientists. Another important finding was that the dynamics of the model were highly sensitive to the asymmetry in the parameters of the excitatory and inhibitory neuronal populations.

An aspect not considered in the Moyse model was the presence of astrocytes that release inhibitory gliotransmitters and, thus, hyperpolarize target neurons. While experimental evidence on the existence of hyperpolarizing astrocytes is relatively negligible, there is specific evidence nonetheless (21, 22). We introduced a hyperpolarizing astrocyte population in their model intending to make it more biologically relevant and explore its behavior to generate further experimentally testable hypotheses that progress research on inhibitory gliotransmitters and hyperpolarizing astrocytes.

Due to the scant data available on these cellular entities, the problem of setting parameters that optimally abstract their biological behavior posed a conundrum. As a result, we employed the Elementary Effects (EE) test, a supplemental strategy for setting parameters wherein the sensitivity of the model to various parameters was tested. Although such analyses have been used extensively in environmental modeling, their potential for answering biological questions has not been fully unearthed in computational neuroscience (23, 24).

## 2 Methodology

In this study, we developed a rate model to investigate the emergence of Up-Down dynamics in a neural network composed of an excitatory population *E*, an inhibitory population *I*, a depolarizing astrocyte population *A*_*d*_, and a hyperpolarizing population, *A*_*h*_. Fig 1 shows the connections. The excitatory population has an adaptation mechanism *a*, which applies an additive hyperpolarizing current to population *E* and grows proportionally to its firing rate. As a result, adaptation is the primary intrinsic mechanism for the emergence of Up-Down dynamics in the model. Each population, however, receives a fluctuating external input. The model’s firing rate dynamics are given by:

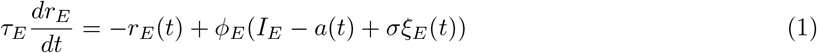

and

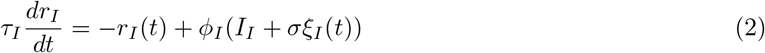

where *r*_*X*_ is the firing rate of population *X* = {*E, I*, *}* and *I*_*X*_ are its recurrent inputs (from the system’s populations). Experiments have shown that oscillatory inputs from the thalamus or other subcortical areas strongly influence cortical Up-Down dynamics. As an independent Ornstein-Uhlenbeck process, the external input *ξ*_*X*_ emulates such an oscillatory external input. *τ*_*x*_ is the population *X* ‘s time constant, and its transfer function is:

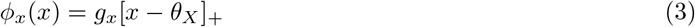

[*z*]^+^ = *z* if *z >* 0 and 0 otherwise, with rectification. The following equation describes the dynamics of the adaptation current *a*(*t*):

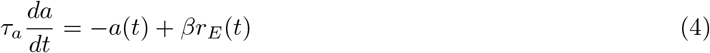

The model proposed in (19) is defined by Eqs (1) to (4). We extended it to account for the influence of both depolarizing and hyperpolarizing astrocytes on the network (Fig 1).

**Figure 1:**
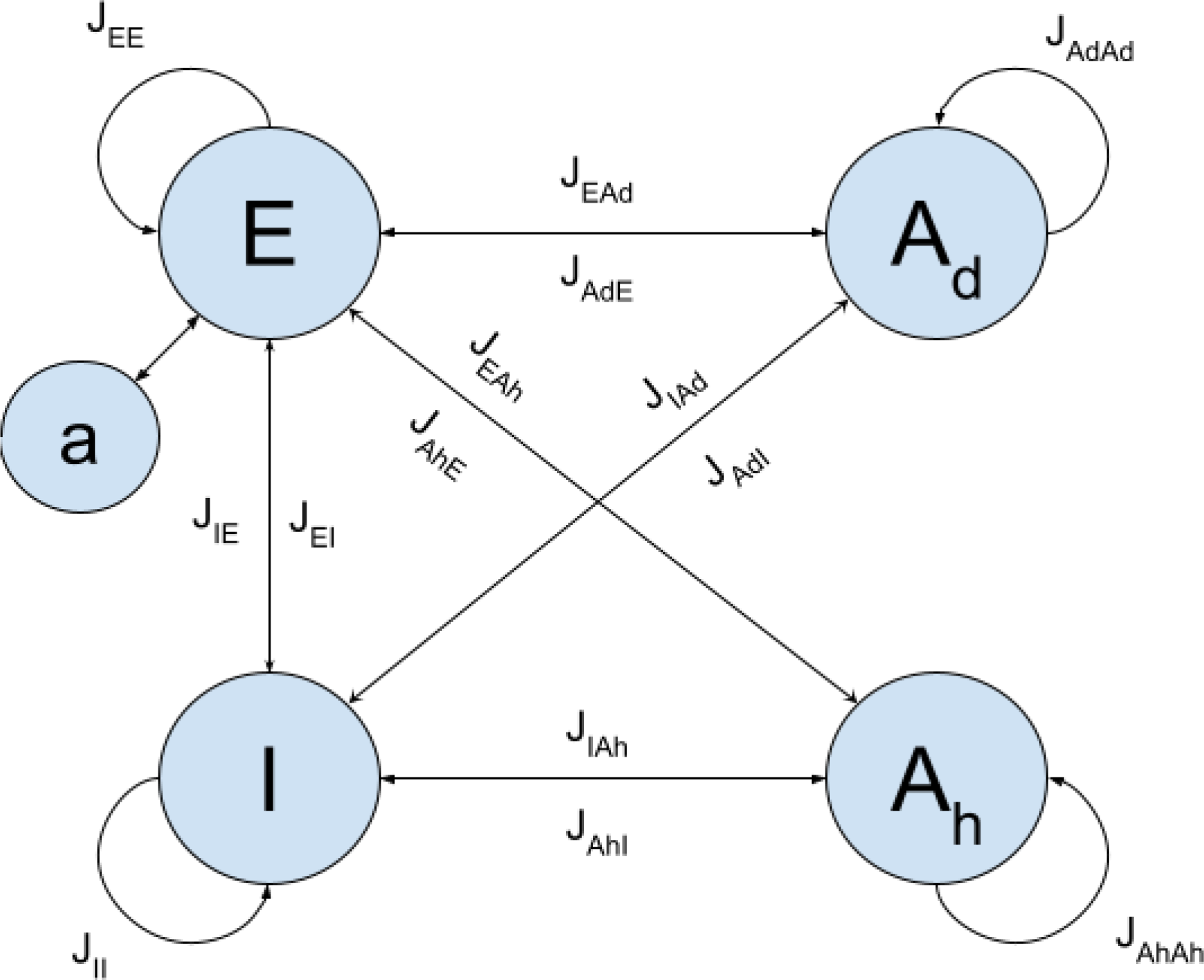
Cell populations in the model and their interactions. E: excitatory neurons, I: inhibitory neurons, *A*_*d*_: depolarizing astrocytes, *A*_*h*_: hyperpolarizing astrocytes. a represents the adaptation mechanism of the excitatory neurons. Arrows represent the interactions, quantified by the synaptic strength (*J*). *J*_*XY*_ indicates the synaptic strength from population Y to population X.

At their membranes, astrocytes express various receptors that bind neurotransmitters or neuromodulators released by presynaptic elements at the tripartite synapse, such as glutamate, GABA, acetylcholine, or taurine. Neuronal activity is integrated inside the astrocyte via these receptors, resulting in a complex signal of astrocytic intracellular *Ca*^2+^. In response to this *Ca*^2+^ transient, astrocytes can, under certain conditions, release gliotransmitters in the synapse. Gliotransmitters can hyperpolarize or depolarize the neuronal membrane after binding to the pre-or post-synaptic element of the tripartite synapse.

This theory of neuron-astrocyte interactions proposes that the astrocytic response to presynaptic neuronal activity is analogous to the neuronal integration process. Incorporated as a calcium trace in astrocytes, the presynaptic neuronal activity causes a peak-like release of gliotransmitters, which in turn affects the voltage of the postsynaptic membrane. The main differences are that gliotransmitter release dynamics and integration timescales in astrocytes differ from electrical signaling in neurons. Astrocytic *Ca*^2+^ does not appear to have an equivalent to inhibitory/hyperpolarizing neuronal inputs that decrease membrane potential. We, therefore, chose to use a similar form of rate equations as Eqs (1) or (2) to model astrocyte activity, but with different time scales. The rate of gliotransmitter release by the astrocyte is expressed as:

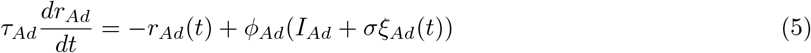

and

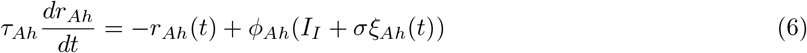

where *τ*_*Ad*_, *τ*_*Ah*_ *>> τ*_*I*_ and *τ*_*Ad*_, *τ*_*Ah*_ *>> τ*_*E*_. Since there is no proof that external inputs are restricted to a particular type of brain cells, an external oscillatory input was added to the astrocyte populations, as well.

We can, thus, define the recurrent inputs as follows:

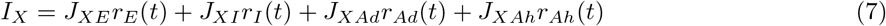

with now *X* = {*E, I, A*_*d*_, *A*_*h*_}. The synaptic couplings *J*_*XY*_ (with *X, Y* = {*E, I, A*_*d*_, *A*_*h*_}), describe the strength of the connection from population *Y* to *X*. For the astrocyte populations, *A*_*d*_ and *A*_*h*_, the right-hand side of Eq (7) contains 3 terms instead of 4 (*A*_*d*_ does not directly influence the rate of gliotransmitter release of *A*_*h*_, and vice versa). Also, note that both *E* and *I* increase the rate of gliotransmitter release in astrocytes.

### 2.1 Fixed Points and Linear Stability Analysis

We followed a mathematical procedure similar to the one described in (19) and (20).

### Noiseless Model

We begin with the rate model specified by Eqs (1) to (6) and ignore the external noisy input first. In this scenario, the system’s nullclines are given by:

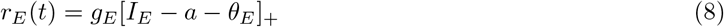

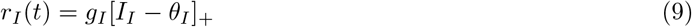

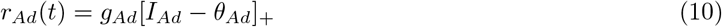

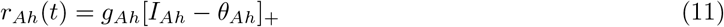

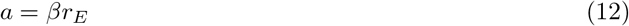

The values of the rates at a fixed point cannot be negative due to the rectification functions in Eqs (8) to (12). Because the rate model equations (1) to (6) are piecewise-smooth systems, a rigorous investigation of the stability of their fixed points would need specialized analysis methods (28). We leave this analysis for now and suppose that all fixed points remain distant from the switching manifolds where the arguments of the rectification functions change signs, and then proceed to linear stability analysis in each area.

A Down state is a fixed-point in which both neuronal populations are mute, i.e. *r*_*E*_ = *r*_*I*_ = 0 spks/s. According to Eq (12), adaption *a* likewise vanishes in such a Down state. For the Down fixed-point to exist, the rectification functions of Eqs (8) and (9) impose *θ*_*E*_ ≥0 and *θ*_*I*_ ≥ 0. Moreover, *θ*_*E*_ *<* 0 would imply that *r*_*Ad*_ *<* 0 and *r*_*Ah*_ *<* 0 at the fixed-point (because *r*_*E*_ = 0 spks/s), which is incompatible with Eqs (10) and (11), respectively. As a result, the Down state exists only if *θ*_*E*_ *≥* 0 and θ_*I*_ *≥* 0. If, in addition, θ_*Ad*_ *≥* 0 and *θ*_*Ah*_ ≥ 0, the Down state is (*r*_*E*_, *r*_*I*_, *r*_*Ad*_, *r*_*Ah*_, *a*) = (0, 0, 0, 0, 0). Assuming that all of the rectification functions in Eqs (1), (2), (5), and (6) disappear in the vicinity of the fixed point, linear stability analysis shows that this Down state is stable (eigenvalues of the Jacobian: {−1*/τ*_*E*_, *−*1*/τ*_*I*_, *−*1*/τ*_*Ad*_, *−*1*/τ*_*Ah*_, *−*1*/τ*_*a*_}).

In the case of *θ*_*Ad*_ < 0 and *θ*_*Ah*_ < 0, still with *θ*_*E*_ ≥ 0 and *θ*_*I*_ ≥ 0, we suppose that the arguments of the rectification function in Eqs (5) and (6) are absolutely positive, whereas the rectification functions in Eqs (1) and (2) disappear. The nullclines for *r*_*Ad*_ and *r*_*Ah*_, Eqs (10) and (11), are then *r*_*Ad*_ = *g*_*Ad*_(*J*_*AdAd*_*r*_*Ad*_ *− θ*_*Ad*_) and *r*_*Ah*_ = *g*_*Ah*_(*J*_*AhAh*_*r*_*Ah*_ *− θ*_*Ah*_). As a result, there is still a positive Down fixed-point (*r*_*E*_, *r*_*I*_, *r*_*Ad*_, *r*_*Ah*_, *a*) = (0, 0, *−g*_*Ad*_*θ*_*Ad*_*/*(1 *− g*_*Ad*_*J*_*AdAd*_), *−g*_*Ah*_*θ*_*Ah*_*/*(1 *− g*_*Ah*_*J*_*AhAh*_), 0) but only for:

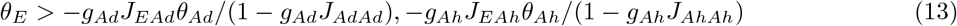

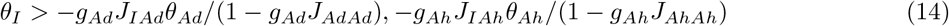

and

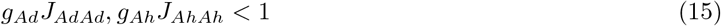

Close to this fixed-point, the Jacobian matrix reads

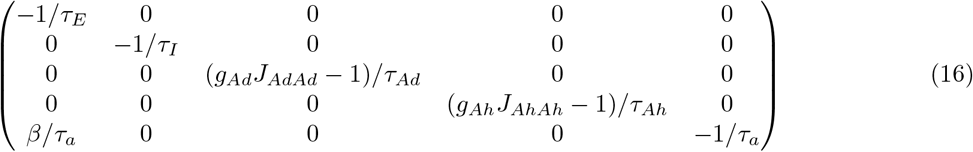

As a result, stability is guaranteed whenever *g*_*Ad*_*J*_*AdAd*_ *<* 1 and *g*_*Ah*_*J*_*AhAh*_ *<* 1, i.e. when the equilibrium values for *r*_*Ad*_ and *r*_*Ah*_ exist.

We chose to omit stability analyses for the scenario where *θ*_*Ad*_ and *θ*_*Ah*_ have opposite signs, considering that our proposed model contains similar reference values for these two parameters.

To identify an Up state fixed-point with non-zero rates, we follow (20) and replace the value of the adaptation at equilibrium, *a* = *β*_*rE*_, assuming that the arguments of all the rectification functions in Eqs (1), (2), (5), and (6) are all strictly positive. This results in:

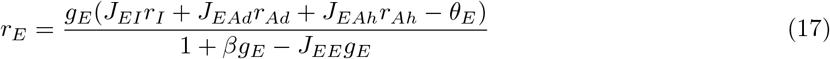

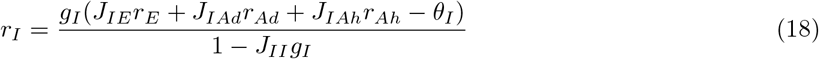

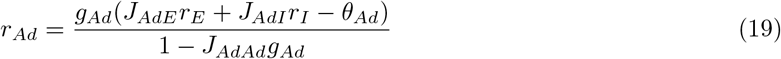

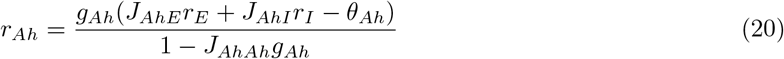

A prerequisite for the existence of the Up state fixed-point is that the right-hand side of Eqs (17), (18), (19), and (20) be positive. Given our reference parameters (Table 1), the most restrictive condition is the condition on *r*_*I*_, i.e., *r*_*I*_ *>* 0. Furthermore, using our reference parameters, another requirement for the presence of the Up state fixed-point becomes

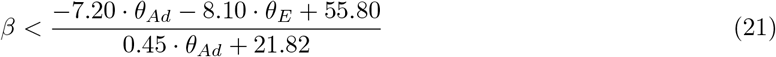

The complexity of the analytical expression of Eq (21) limited its utility and therefore, has been omitted. Considering that in the neighborhood of the Up fixed-point, all the rectification functions of the model are positive, the Jacobian matrix reads

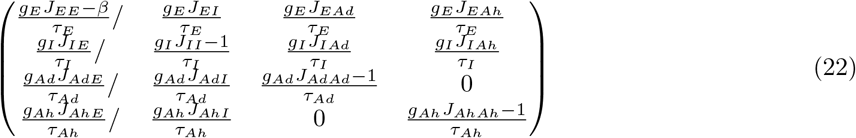

Analytical equations for the eigenvalues of this matrix can be obtained, but they are far too complex to be useful. We, therefore, estimated their values numerically to explore the stability of the Up fixed-point.

**Table 1:** Parameters for the rate model. The list of parameters derived from our parameter optimization strategy and employed for simulating the model.

| Parameter | Value | Definition | Parameter | Value | Definition |
| --- | --- | --- | --- | --- | --- |
| $\tau_E$ | 10 ms | time constant, E | $J_{EE}$ | 5 s | strength, E→E |
| $\tau_I$ | 2 ms | time constant, I | $J_{EI}$ | -1 s | strength, I→E |
| $\tau_{Ad}$ | 20 ms | time constant, A<sub>d</sub> | $J_{EAd}$ | 1 s | strength, A<sub>d</sub>→E |
| $\tau_{Ah}$ | 20 ms | time constant, A<sub>h</sub> | $J_{EAh}$ | -1 s | strength, A<sub>h</sub>→E |
| $\theta_E$ | 7.5 | threshold, E | $J_{II}$ | -0.5 s | strength, I→I |
| $\theta_I$ | 25 | threshold, I | $J_{IE}$ | 10 s | strength, E→I |
| $\theta_{Ad}$ | -9 | threshold, A<sub>d</sub> | $J_{IAd}$ | 0.5 s | strength, A<sub>d</sub>→I |
| $\theta_{Ah}$ | -3.5 | threshold, A<sub>h</sub> | $J_{IAh}$ | -0.5 s | strength, A<sub>h</sub>→I |
| $g_E$ | 1 Hz | gain, E | $J_{AdAd}$ | 0.1 s | strength, A<sub>d</sub>→A<sub>d</sub> |
| $g_I$ | 4 Hz | gain, I | $J_{AdE}$ | 0.5 s | strength, E→A<sub>d</sub> |
| $g_{Ad}$ | 1 Hz | gain, A<sub>d</sub> | $J_{AdI}$ | 0.5 s | strength, I→A<sub>d</sub> |
| $g_{Ah}$ | 1 Hz | gain, A<sub>h</sub> | $J_{AhAh}$ | 0.1 s | strength, A<sub>h</sub>→A<sub>h</sub> |
| $\tau_a$ | 500 ms | time constant, a | $J_{AhE}$ | 0.5 s | strength, E→A<sub>h</sub> |
| $\beta$ | 3.6 | strength, a | $J_{AhI}$ | 0.5 s | strength, I→A<sub>h</sub> |
| $\sigma$ | $3.5\sqrt{2}$ | noise standard | | | |

### The effect of noise on the model

The inclusion of noise, on the other hand, results in a situation where spontaneous Up-Down transitions can occur even in regions where the Down state is stable and the Up state is unstable, through a mechanism wherein noise triggers the Down to Up switches and adaptation prompts the reverse Up to Down transitions. In (20) it is argued that this subregion begins when the Up state is unstable due to adaptation alone, a requirement that can be inferred from Eq (21) with *β* = 0:

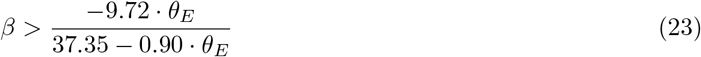

Similarly, a symmetrical regime exists in the area where the Up state is stable and the Down state is unstable, with noise triggering Up-to-Down transitions and adaptation triggering Down-to-Up shifts. According to (20), this area is defined by the circumstance in which adaptation in the Up state is substantial enough to offset the influence of *θ*_*E*_, i.e., *βr*_*E*_ + *θ*_*E*_ *>* 0, where *r*_*E*_ is provided by Eq (16), which produces:

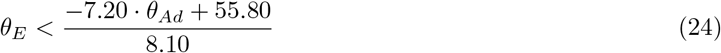

The complexity of the analytical forms of Eqs (23) and (24) limited their utility and therefore, have been omitted.

### 2.2 Parameter Estimation

The experimental data available for estimating the rate model’s parameters are highly variable. For example, the values used to quantify the experimental measurements in (20) change dramatically from one repetition of the experiment to the next. The diversity would most likely be considerably greater with other researchers using various experimental settings and recording on different brain areas. As a result, traditional parameter estimation approaches can only fit one unique repetition of a certain experiment. Still, our goal here is to acquire a more generic perspective of this system. Thus, we opted for a parameter estimation method similar to the one described in (19) and used the reference values that were estimated from experimental literature in their study. To avoid oversimplification, we then determined the relative sensitivity of the model dynamics to various parameters (see below). From this Sensitivity Analysis (SA), a set of three parameters was chosen (*θ*_*E*_, *θ*_*Ad*_, and *β*) that most strongly influenced the rate model dynamics. The SA was followed by bifurcation studies to select a set of optimal values that resulted in reasonable similarity to the experimental data provided in (20).

### 2.3 Sensitivity Analysis

Previously, SA has been employed to assess the qualitative and quantitative dependency of model parameters on model behavior (25). We used the EE test or Morris method, a type of SA, to find the parameters that had the most profound impact on the dynamics of the model (26). The Morris method follows the One-At-a-Time (OAT) approach that evaluates the measurement of perturbation in the model output by drawing samples from the spread of the predefined parameter space. A list of *N = 10* parameters was chosen with varying ranges and the elementary effects for the *n*^*th*^ parameter were calculated with Eq (25), where *δ* is the increment:

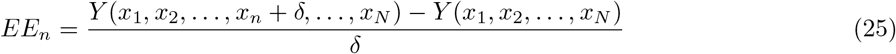

To measure the mean of the elementary effects for any parameter Eq (26) was used, where *n*_*R*_ is the total number of elementary effects for the *n*^*th*^ parameter:

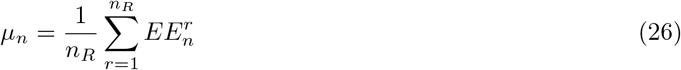

Similarly, standard deviations of the elementary effects were derived with Eq (27):

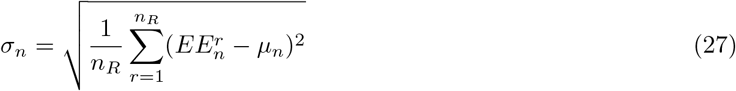

Owing to its low computational processing requirements, the EE test is one of the most often utilized SA screening procedures. It is a viable alternative in computationally expensive models and where other SA approaches, such as variance-based tests, are not practicable. We did SA for ten parameters and characterized the highly sensitive parameters as free parameters compared to the Moyse model. It was implemented using the Sensitivity Analysis For Everyone (SAFE) toolkit, an open-source program (23).

## 3 Results

The basic numerical simulations in Fig 2 show the dynamics of the rate model. When simulated with the same set of parameters as Jercog, Roxin, et al., 2017, the Moyse model exhibited bistability and spontaneous Up-Down state switching (19, 20). Note that when the parameters were optimized to fit experimental data, the Jercog model failed to exhibit bistability, demonstrating the essentiality of a third excitatory cell population for bistability. The model perturbation by incorporating hyperpolarizing gliotransmission underscores its susceptibility to qualitatively different architectures. Nevertheless, a bistable regime was identified in the dynamics continuum, when simulated in a parameter space distinct from the Moyse model.

**Figure 2:**
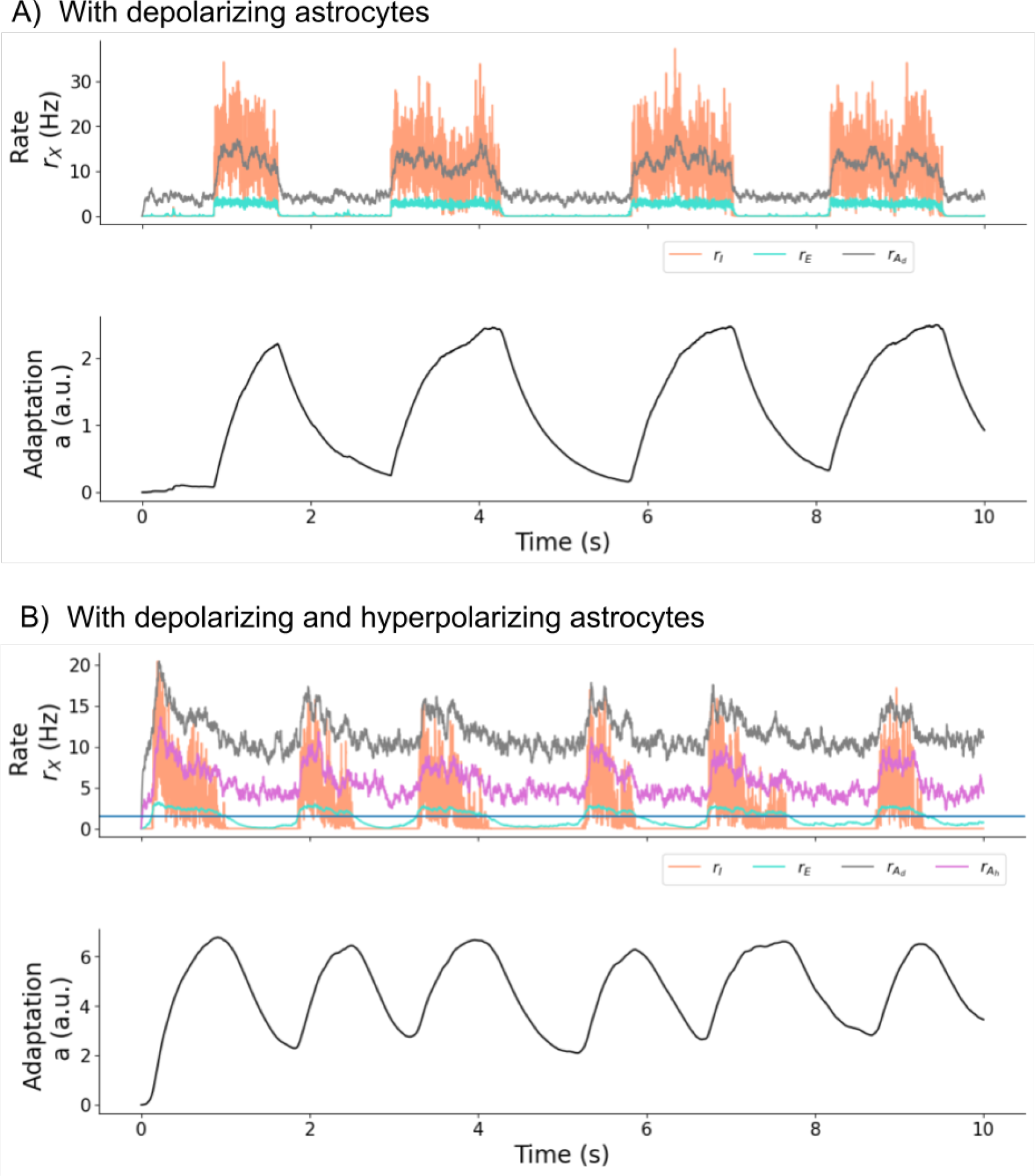
Spontaneous transitions between Up and Down states. (A) The model with only depolarizing astrocytes (*J*_*XAh*_ = *J*_*AhX*_ = 0) demonstrates cortical Up-Down state switching. (B) With the addition of a hyperpolarizing astrocytic population and with parameter optimization (see Methodology), the model continues to show Up-Down switching.

To delineate the dynamical regimes in which the model was bistable as well as exhibited spontaneous Up-Down transitions, we performed a stability analysis similar to the one in (19). The resultant Eq (23) and (24), thus, delimited the bistable dynamical regime as shown in Fig 3. Fig 3 specifically demonstrates the dynamical regimes as a function of *β, θ*_*E*_, and *θ*_*Ad*_, which is in concordance with our SA results (see below). Low values of *β* and *θ*_*E*_ are expected to give rise to a regime of persistently high firing rates, i.e., a stable Up state. Large values of *θ*_*E*_, on the other hand, are likely to result in a quiet regime, or Down state, in which neuronal firing rates disappear. The system is expected to switch spontaneously between periods of high population rates and periods of total silence, i.e., U↔ D dynamics, between these two zones.

**Figure 3:**
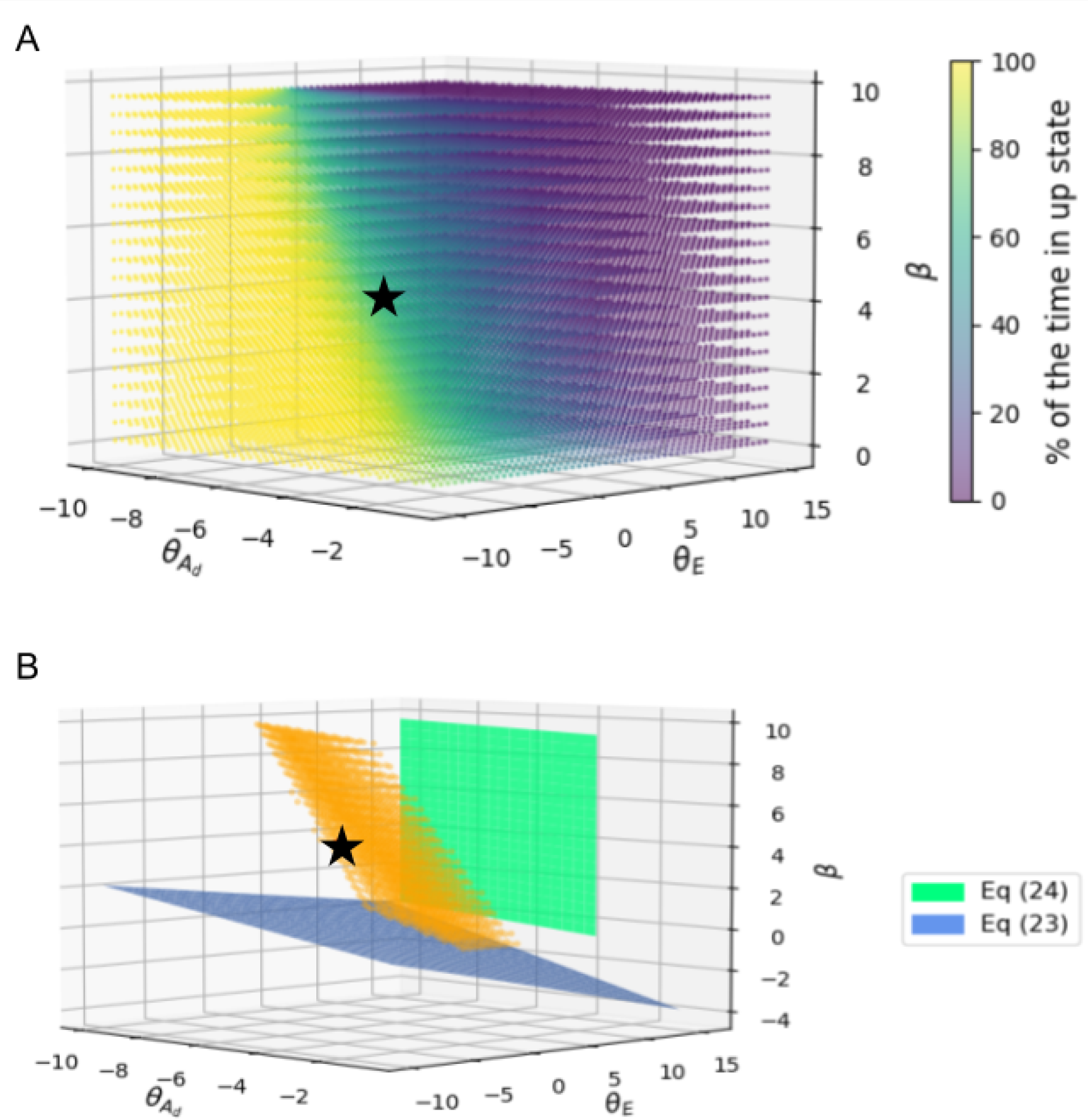
Fixed points and linear stability analysis. (A) The heat map shows the continuum of model dynamics across a range of values of the high-sensitivity parameters. (B) The sliced heat map in orange (% Up in the range 40 to 50%) bounds the frontier regions beyond which the model loses stability and converges to either an Up state fixed-point or a Down state fixed-point. The planes represent Eq (24) and Eq (23) which delineate the stable region estimated by the linear stability analysis (see Methodology). The black star represents the choice of parameters shown in Table 1.

The results from the EE test are shown in Fig 4. The model dynamics are highly sensitive to changes in *β, θ*_*E*_, and *θ*_*Ad*_, as compared to other parameters. It is to be noted that the presence of gliotransmission couplings pushes the model dynamics from the D region to the U*←→*D region with the same set of values for *β* and *θ*_*E*_ (19). The addition of hyperpolarizing gliotransmission seems to perturb model dynamics, although *J* is a low-sensitivity parameter (data not shown). In contrast, when exploring a larger parameter space that includes values of *J* becoming zero or changing sign, it becomes a high-sensitivity parameter owing to qualitative changes in the model architecture. The addition of a hyperpolarizing astrocyte population did not have a significant effect on the ability of the model to spontaneously transition between Up and Down states and exhibit bistability. The presence of hyperpolarizing gliotransmission made the parameter space for *β* and *θ*_*E*_ in which the model is bistable relatively narrower when compared with the Moyse model, probably due to its inhibitory effects on the model dynamics.

**Figure 4:**
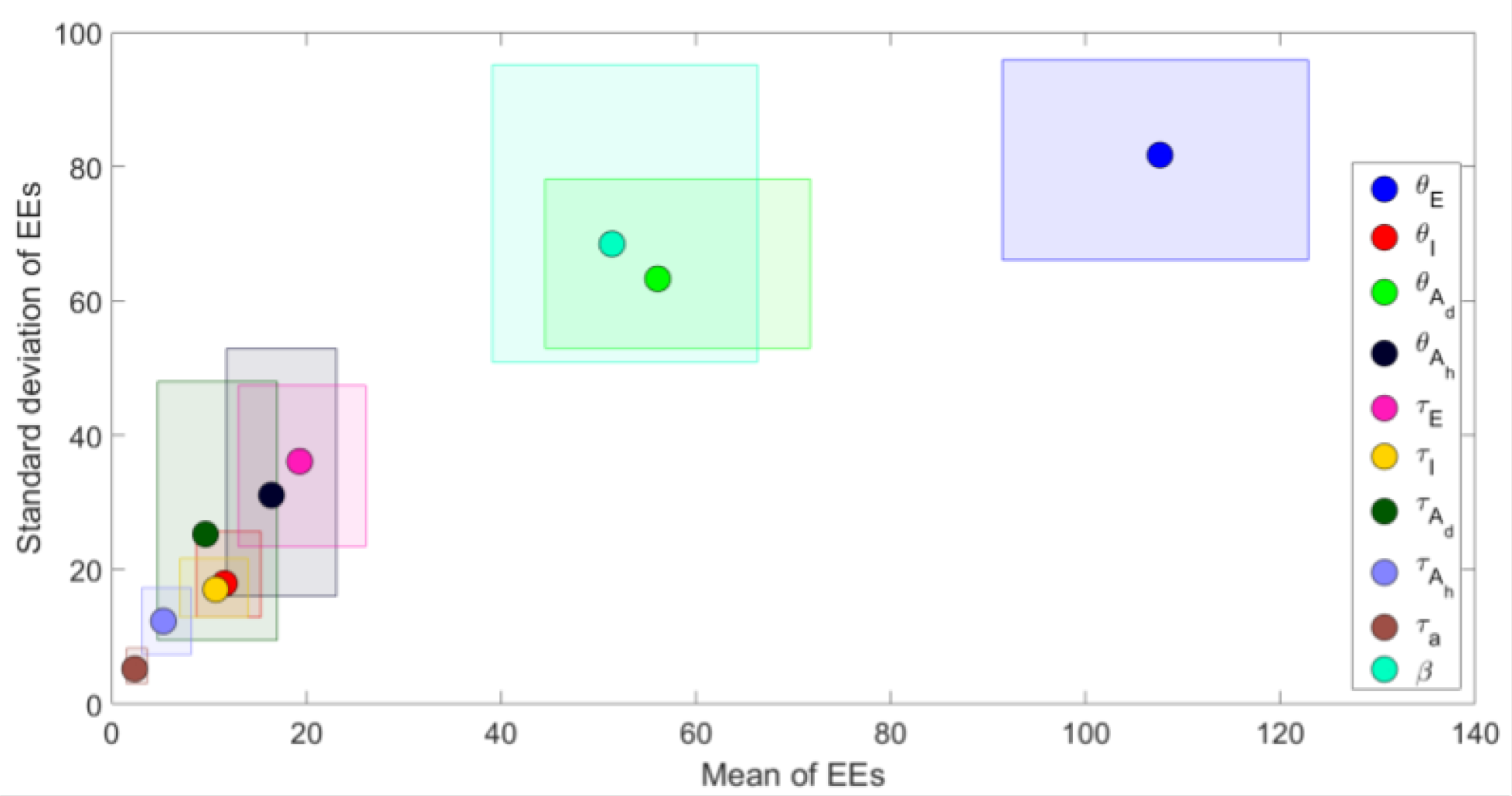
Differential sensitivity of the model dynamics to various parameters. The means of the sensitivity of the model to various parameters is plotted against the standard deviation of EEs with confidence intervals derived from bootstrapping. Parameters are ranked for influence based on their relative position. EEs: Elementary Effects.

## 4 Discussion

Our theoretical work demonstrates that neuronal systems with hyperpolarizing gliotransmission can be mathematically stable. This suggests that hyperpolarizing gliotransmission may play a role in the development of Up-Down dynamics. An important implication of these findings is that this hypothesis could be tested experimentally using electrophysiology and calcium imaging methods. Although evidence is limited, it appears that hyperpolarizing astrocytes play an important role in the refinement of neuron-astrocyte circuitry. Hyperpolarizing astrocytes, like interneurons, can contribute to the creation of complex information processing architectures as well as the modification of existing architectures, resulting in increased efficiency (27). We speculate that hyperpolarizing astrocytes could contribute to functional neuroscience outcomes in numerous ways in the context of the available evidence. The concurrent development of theoretical models that explore their roles to guide and validate experimental findings is the broader goal that this paper serves.

We represented astrocytes computationally in the same manner that Moyse and Berry, 2022 did, with astrocytes and neurons having identical qualitative properties but different parameter spaces. However, there is a compelling case to be made for the growing number of astrocyte and neuron-astrocyte synapse models that recognize the more complex and variable roles of astrocytes, as opposed to the simplistic view of astrocytes as supportive connective tissue cells. Aside from gliotransmission, evidence supports their role in plasticity, neuroprotection, information processing, and metabolic interactions with neurons (28). All of these factors may directly or indirectly influence neuronal behavior in the system under consideration, and thus, the functional outcome. As a result, without a comprehensive and holistic understanding of astrocytes, unified modeling of cortical dynamics is rendered incomplete.

Model parameter uncertainty constrained by biology, experimental conditions, and phenomenological abstractions stipulates the use of SA methods such as global sensitivity analysis and the EE test (30). To mitigate the effects of phenomenological abstractions, we used the EE test to determine the optimal model parameters in this paper. Because Moyse and Berry, 2022 used an ad hoc method to estimate parameters, it is assumed that neither could align biologically well. Nonetheless, the behavior of the model appeared to match the experimental evidence presented in (20). Using the EE test, we discovered that the strength of coupling between cell populations is a low-sensitivity parameter in our model, possibly due to neuroplastic changes, further aligning the model biologically. The adaptation strength is a high-sensitivity parameter that is critical to the emergence of Up-Down cortical switching. The excitatory population thresholds (*θ*_*E*_ and *θ*_*Ad*_) are found to influence the model dynamics more prominently than the inhibitory population thresholds (*θ*_*I*_ and *θ*_*Ah*_). This is most likely due to the network being more sensitive to excitation than inhibition due to the low neuronal firing rate when viewed in the context of the biological spectrum (31).

We extended the Moyse rate model of cortical Up-Down state switching in this study to investigate the impact of a hyperpolarizing astrocyte population on model dynamics. While Moyse and Berry, 2022 also developed a spiking model to provide a theoretical framework for the data presented in Jercog, Roxin, et al., 2017, we have omitted the simulations for a spiking model and the associated analytical procedures due to computational constraints. We used similar equations as (19) to model neuronal and astrocyte populations and their interconnections, while retaining the properties of spontaneous Up-Down state switching and bistability. We supplemented an ad hoc method of parameter estimation with the EE test, a type of SA that identifies the parameters that have the greatest impact on model behavior. From a mathematical standpoint, the SA findings corroborated experimental findings, demonstrating that the threshold of excitatory cellular populations and the strength of adaptation strongly influence model dynamics while synaptic strengths do not, as long as no qualitative changes in model architecture are considered. With the application of SA methods, we demonstrated its use in computational neuroscience, where its potential remains untapped. Our findings suggest that hyperpolarizing gliotransmission may be well involved in the development of cortical Up-Down state switching. We hypothesize that hyperpolarizing astrocytic populations might be involved in neural information processing. We also anticipate that incorporating hyperpolarizing astrocytic populations will aid in the development of better machine-learning models (32, 33).

## 5 Acknowledgements

We are grateful to Dr. Hugues Berry and Lisa Blum Moyse (University of Lyon, Villeurbanne, France) for their invaluable assistance in elucidating the mathematics of the linear stability analysis and providing guidance on the computational code for our rate model. We also wish to express our appreciation to Aiswarya P S (IISER Thiruvananthapuram, Thiruvananthapuram, India) for her invaluable insights into the calculations for the linear stability analysis.

## Notes

### Competing Interest Statement

The authors have declared no competing interest.

